# Hippocampal contextualization of social rewards

**DOI:** 10.1101/2023.05.12.540475

**Authors:** Joana Mendes Duarte, Kaizhen Li, Robin Nguyen, Stéphane Ciocchi

## Abstract

Acquiring and exploiting memories of rewarding experiences is critical for survival. The spatial environment in which a rewarding stimulus is encountered regulates memory retrieval. The ventral hippocampus (vH) has been implicated in contextual memories involving rewarding stimuli such as food, social cues or drugs. Yet, the spatial representations and circuits underlying contextual memories of socially rewarding stimuli are poorly understood. Here, using *in vivo* electrophysiological recordings during a social reward contextual conditioning paradigm in mice, we showed that vH neurons discriminate between contexts with neutral or acquired social reward value and exhibit a preferential remapping of their place fields to the context previously paired with social reward cues. The formation of context-discriminating vH neurons following learning was contingent upon the presence of salient reinforcers. Moreover, vH neurons showed different contextual representations during retrieval of social reward and fear contextual memories, suggesting different vH circuits underlie positively and negatively valenced contextual memories. Finally, optogenetic inhibition of locus coeruleus (LC) projections in the vH selectively disrupted social reward contextual memory by impairing vH contextual representations. Collectively, our findings reveal that the vH integrates contextual and social reward information, with memory encoding of these representations supported by input from the LC.

## Introduction

The survival of animals depends on their ability to access and use memories of rewarding and aversive experiences (LeDoux, 2000). Memory-guided behaviour depends on the brain’s capacity to simultaneously construct internal representations of salient emotional experiences and spatial environments in which these experiences occur. In particular, learning about social cues is highly adaptive as it enables individuals to locate predators, resource competitors and reproductive mates (Chen and Hong, 2018). Certain social cues can act as unconditioned stimuli that motivate and reinforce behaviour. For instance, male and female rodents learn to approach and preferentially occupy environments (i.e. contexts) that had previously contained cues of the opposite sex (Beny-Shefer et al., 2017; Chen and Hong, 2018; Dai et al., 2022). While the neuronal mechanisms involved in the processing of social cues have been largely investigated (Bergan et al., 2014; Gunaydin et al., 2014; Levy et al., 2019), the neuronal circuits and representations underlying contextual memories related to social cues remain poorly understood.

The vH is a brain region involved in encoding and retrieving affective memories (Fanselow and Dong, 2010) and mediates behavioural responses associated with positive and negative states (Bannerman et al., 2003; Britt et al., 2012; Ito et al., 2008; LeGates et al., 2018; Okuyama et al., 2016; Xu et al., 2016; Zhou et al., 2019). Specifically, vH neurons respond to affective zones within a behavioural arena and exhibit large place fields that map onto contexts (Jung et al., 1994; Keinath et al., 2014; Komorowski et al., 2013). Furthermore, disrupting vH activity in learning tasks with positive and negative unconditioned stimuli impairs memory, such as during contextual reward conditioning (LeGates et al., 2018; Zhou et al., 2019), social recognition (Okuyama et al., 2016; Meira et al., 2018; Phillips et al., 2019) and contextual fear conditioning (Maren and Holt 2004, Jimenez et al., 2017; Xu et al., 2016). Moreover, neuronal populations within the vH are known to send efferents to- and receive afferents from a variety of brain regions involved in innate and conditioned behaviours (Gergues et al., 2020; Wee and MacAskill, 2020). Together, these findings implicate the vH as a core structure mediating context-dependent emotional behaviours receiving convergent spatial and salience information.

The vH receives neuromodulatory inputs from several brain regions that may be involved in regulating synaptic plasticity mechanisms underlying learning (Palacios-Filardo and Mellor, 2019). Of note, the locus coeruleus (LC) is recruited by salience or novelty (Sara et al., 2009; Takeuchi et al., 2016), and is a major source of both noradrenaline and dopamine in the hippocampus (Kempadoo et al., 2016; Takeuchi et al., 2016; Lemon et al., 2009; Smith and Greene, 2012). Noradrenergic signaling enhances CA1 long-term potentiation (Lin et al., 2003; Liu et al., 2017; Qian et al., 2012; Thomas et al., 1996; Wang et al., 2010), and LC inputs to the dorsal hippocampus facilitate spatial and contextual learning (Kempadoo et al., 2016; Takeuchi et al., 2016;), support the enrichment of place cells near reward zones (Wagatsuma et al., 2018) and the linking of temporally distinct contextual memories (Chowdhury et al., 2022), including place-reward learning (Kaufman et al., 2020). The function of LC projections to the vH, on the other hand, has not yet been explored.

In the present study, we examined the activity of vH neurons using *in vivo* single-unit electrophysiological recordings and calcium imaging combined with optogenetic manipulations of the LC in mice during a social reward contextual memory paradigm. Our findings indicate that the vH is critical for associating contexts with social rewards via a process dependent on LC activity.

## Results

### The vH discriminates contexts following social reward contextual conditioning

The vH has been shown to encode contexts associated with natural rewards (Ciocchi et al., 2015; Riaz et al., 2017), including social rewards (Ciocchi et al., 2015, Riaz et al., 2017, LeGates 2019). To identify the neuronal correlates of contextual memory for social reward in the vH, we subjected freely-moving mice (3-5 months old, *n* = 23, C57BL6/J) to a conditioned place preference task (CPP) using social-odour reward while performing chronic single-unit recordings (Fig. 1A and Supplementary Fig. 1). Mice were first habituated (Pre-Train Test) to a maze with two chambers with different features (i.e. contexts) connected by a central alley. Then, throughout three consecutive training sessions separated by one hour, mice were placed in the neutral context containing a neutral odour (clean rodent bedding mixed with 1% cinnamon) and a social reward context containing female odour (i.e. home-cage bedding of adult females naturally impregnated with female odours and pheromones) which acted as a conditioned reinforcer. Fourteen hours following training, mice were re-exposed to the maze – in the absence of added odours – to test social reward CPP memory (Post-Train Test). Memory strength was measured by the degree of preference for the social reward context which was calculated as a discrimination score using the time spent in each context of the maze (i.e. neutral vs. social reward) (Fig. 1B). During training, mice exhibited a strong preference for the context and zone paired with social reward (Fig. 1C and Supplementary Fig. 2A). This preference was maintained during the Post-Train Test session and was elevated compared to the Pre-Train Test session (Fig. 1C and Supplementary Fig. 2A), indicating that mice formed a memory following social reward contextual conditioning.

**Figure 1.**
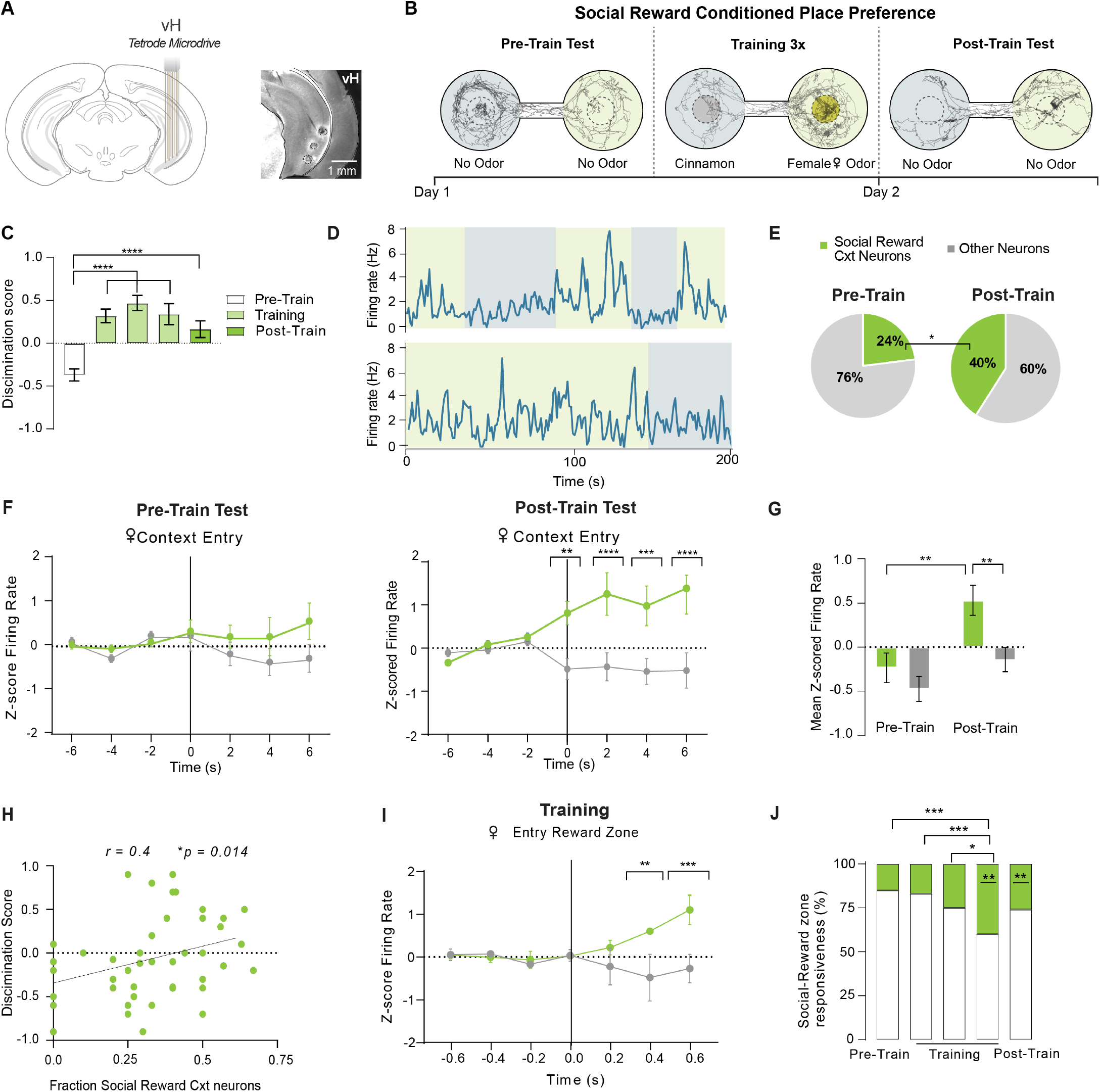
The vH discriminates contexts following social reward contextual conditioning. **(A)** Left, schema of the chronic implantation of tetrodes in the vH using a tetrode microdrive. Right, representative image of recording sites in the vH after electrolytic lesions. **(B)** Experimental design of the social reward CPP paradigm with representative mouse trajectories during the Pre-Train, Training, and Post-Train Test sessions. **(C)** Discrimination score during social reward CPP. Repeated measures one-way ANOVA *F*_(2.801,61.62)_ = 18.48, *P* < 0.0001, *n* = 23 mice. **(D)** Representative example of two social reward context neurons during Post-Train Test session with the corresponding entries in the social reward context (green) and neutral context (grey). **(E)** Comparison of social reward context neurons proportions between the Pre-Train Test and Post-Train Test sessions. Chi-square between groups, *P* = 0.0153, *n* = 23 mice. **(F)** Averaged normalized firing rate (z-score) of social reward context neurons upon entry into each context during the Pre-Train Test (left) and Post-Train Test (right) sessions. Repeated measures two-way ANOVA between groups, Time x Context effect, Left, *F*_(6,768)_ = 1.188, *P* = 0.3106, *n* = 65 neurons, Right, Time x Context *F*_(6,438)_ = 5.940, *P* <0.0001, Context effect *F*_(1,73)_ = 15.65, *P* = 0.0002, Multiple Comparisons between time points, *P* = 0.0022 (0 seconds), *P* <0.0001 (2 seconds), *P* = 0.0006 (4 seconds), *P* <0.0001 (6 seconds), n = 74 neurons, *n* = 23 mice. **(G)** Mean normalized activity of social reward context neurons during the Pre-Train Test and Post-Train Test sessions. Two-way repeated measures ANOVA. Session effect, *F*_(1,88)_ = 10.05, *P* = 0.0021, Context effect *F*_(1,88)_ = 11.54, *P* = 0.0010, social reward context Pre-Train vs. Post-Train, *P* = 0.0048; social reward context vs. neutral context at Post-Train session, *P* = 0.0050, *n* = 45 neurons, *n* = 15 mice. **(H)** Linear positive correlation between behavioural social reward CPP expression in each animal and the proportion of social reward context neurons during Pre-Train and Post-Train sessions. Pearson correlation between groups (*r* = 0.4, *P*= 0.0141, *n* = 23 mice). **(I)** Activity of social reward-responsive neurons during training. Two-way repeated measures ANOVA, time x context effect: *F*_(6,468)_ = 4.344, *P* = 0.0003. Multiple comparisons between social reward vs. neutral context at 0.4 seconds *P* = 0.0020 and 0.6 seconds *P* = 0.0003, *n* = 41 neurons, *n* = 11 mice. **(J)** Proportions of social reward context neurons activated in the reward-zone during social reward CPP, *n* = 68 neurons, *n* = 19 mice Chi-square test between groups. Pre-Train Test session vs. Training 3 *P* = 0.0001, Training 1 vs. Training 3 *P* = 0.0005, Training 2 vs. Training 3 *P* = 0.0224. Chi-square test between reward zone responsive cells vs. non-responsive cells during training session 3 *P* = 0.0044, Post-Train Test session *P* = 0.0031.

Next, we compared the firing rates of individual vH neurons when mice occupied the neutral or social reward context at the Post-Train Test session using a bootstrapping approach. We found vH neurons that exhibited higher overall activity in the social reward context than in the neutral context during the Post-Train Test session which we called social reward context neurons (Fig. 1D). Of particular note, the fraction of vH neurons representing the social reward context increased after training (Fig. 1E). Furthermore, social reward context neurons exhibited increased activity upon entering the social reward context at the Post-Train Test session (Fig. 1F), with this context-discriminating activity observed only after training (Fig. 1F,G). Importantly, the firing rates of social reward context neurons were not correlated with the speed of movement (Supplementary Fig. 2B). Moreover, we found that the preference for the social reward context at the behavioural level was correlated with the proportion of social reward context neurons (Fig. 1H), suggesting that vH neuronal activity representing the social reward context may support memory retrieval. This correlation was absent for vH neurons with a preferential activity in the neutral context (Supplementary Fig. 2C). To further confirm that the activity of social reward context neurons carried selective information about the context, we decoded the context identity from the firing rates of individual mice using a linear support vector machine (SVM) classifier. We found that the classification accuracy was higher with a model trained on correct compared to randomized labels (Supplementary Fig. 2D). Additionally, as a consequence of learning, we observed that a subset of social reward context neurons enhanced their firing when mice entered into the social reward zone during the Post-Train Test session(Fig 1I), in parallel with an increase in their proportion (Fig. 1J). Together, these findings indicate that the vH discriminates contexts following social reward contextual conditioning.

### Social reward context neurons remap upon learning

Previous studies have demonstrated that the vH can represent large areas (Kjelstrup et al., 2008) as well as affective zones within contexts (Komorowski et al., 2013) and contains place cells that remap upon the presentation of an aversive odour (Keinath et al., 2014). Thus, we examined alterations in the place fields of social reward context neurons upon social reward CPP learning. During the Pre-Train Test session, a large fraction of social reward context neurons exhibited place field activity peaks in both contexts, while after training, the peak location remapped toward the social reward context (Fig. 2A, B). We asked whether this observation may be attributed to place field remapping among a subset of neurons. Upon examining the stability of place fields between the Pre-Train and Post-Train Test sessions, we found that vH neurons with peak place field activity in the social reward context during the Pre-Train Test session showed higher field stability compared with those that had peaks located in the neutral context or in both contexts, as indicated by the correlation values between place fields in the Pre- and Post-Train sessions (Fig. 2C). These results indicate that the formation of a social reward contextual memory involves both the remapping of vH place fields towards the social reward context and the maintenance of pre-existing place fields within the social reward context.

**Figure 2.**
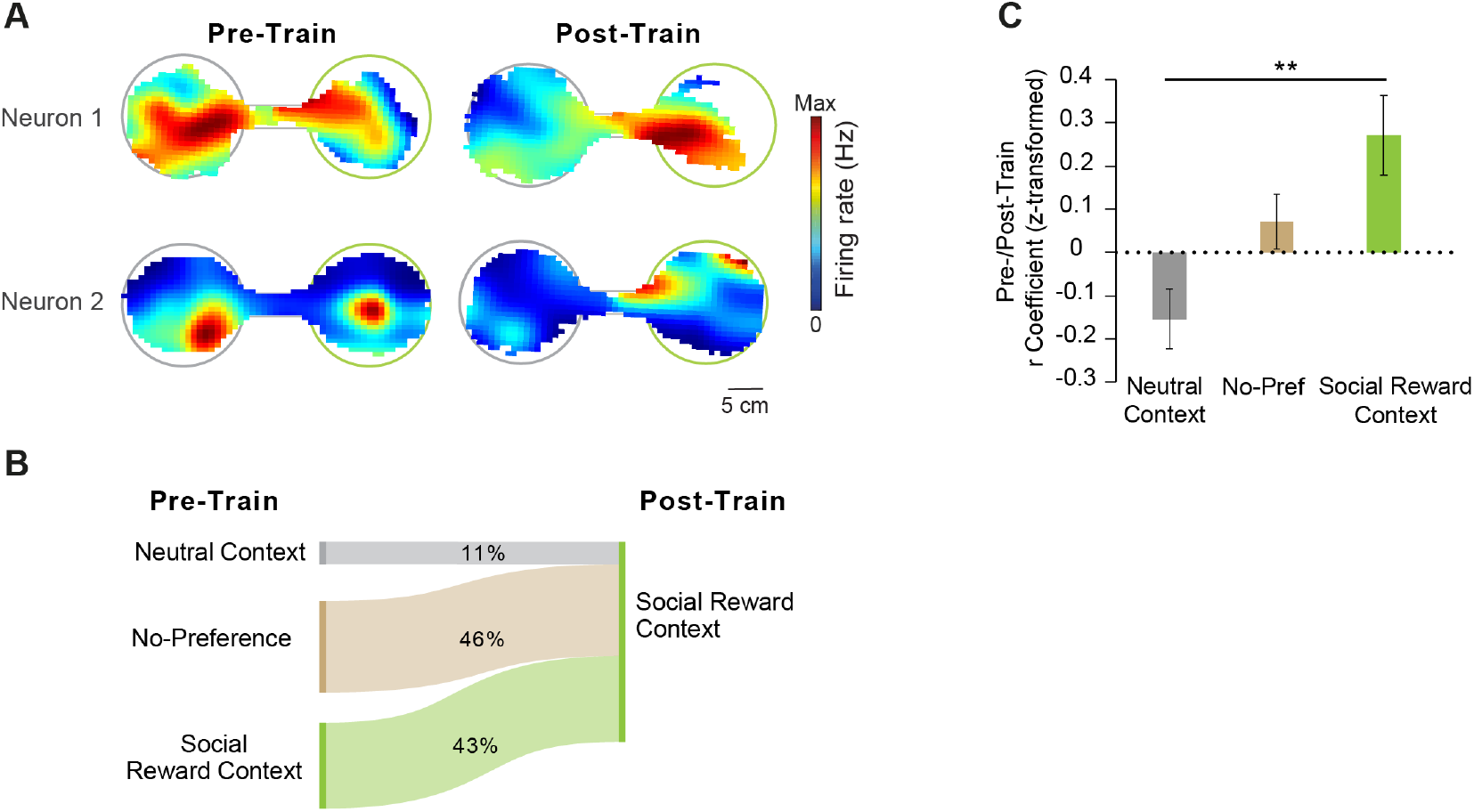
Social reward context neurons remap upon learning. **(A)** Representative examples of the spatial activity of two social reward context neurons that remap during Pre- and Post-Train Test sessions. **(B)** Alluvial flow chart depicting changes in the localization of the peak activity of the place field of the social reward context neurons during Pre-Train Test and Post-Train Test sessions. **(C)** Comparison of normalized r correlations (Fisher’s Z-transformed) of place fields between the Pre-Train Test and Post-Train Test sessions for vH neurons exhibiting the peak of the place field in the social reward context, the neutral context or both contexts during the Pre-Train Test. One-way ANOVA *F*_(2,68)_ = 5.07, *P* =0.0089, n = 74 neurons, *n* = 23 mice.

### Emergence of social reward context neurons is contingent on associative learning

The selective vH contextual activity observed may be tied to an associative emotional experience or reflect a non-associative process such as contextual sensitization or the mere passage of time. To evaluate if the presence of the unconditioned social reward reinforcer was crucial for the development of context-discriminating activity, a control CPP paradigm involving only neutral odours was conducted in a separate cohort of mice (*n* = 5 mice) followed by the classical social reward CPP (Fig. 3A). In the control CPP paradigm, mice showed higher variability in time spent in the two contexts during learning and an absence of preference for a particular context in the Post-Train Test (Fig. 3B, C). This contrasts with behaviour in the social reward CPP paradigm, in which mice successfully learned to form an association between the paired context and the social reward (Fig. 3B, C). Then, we compared the activity of vH social reward context neurons between the control CPP and the social reward CPP paradigm during the Post-Train Test, and observed that they exclusively formed during the social reward CPP but not the control CPP paradigm (Fig. 3D). Thus, both social reward contextual memory and the accompanying vH contextual representations rely on salient reinforcers during associative learning.

**Figure 3.**
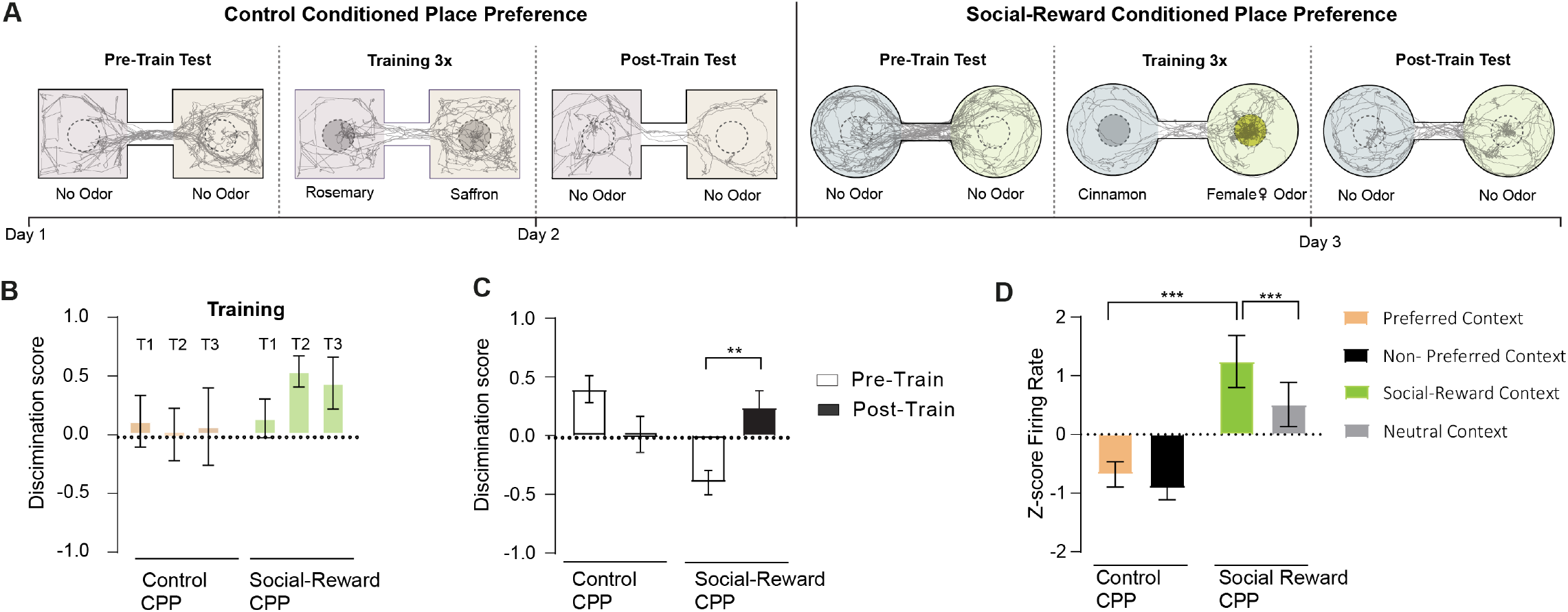
Emergence of social reward context neurons is contingent on associative learning. **(A)** Experimental design of the control CPP and social reward CPP paradigm with representative mouse trajectories during the Pre-Train, Training, and Post-Train Test sessions. **(B)** Discrimination score during the training of the control CPP and social reward CPP, *n* = 5 mice. Two-way repeated measures ANOVA between the groups, task effect, *F*_(1,8)_ = 1.251, *P* = 0.258. T1, T2, T3 correspond to the three consecutive learning sessions. **(C)** Discrimination score during the Pre-Train and the Post-Train Test sessions in the control CPP and social reward CPP. Two-way repeated-measures ANOVA between the groups, session x task interaction effect, *F*_(1,8)_ = 21.31, *P* = 0.0017. Multiple comparisons between Pre-Train and Post-Train Test during social reward conditioning, *P* = 0.0071, *n* = 5 mice. **(D)** Averaged normalized firing rate activity in each context during control CPP and social reward CPP paradigms. Two-way repeated-measures ANOVA between groups, context vs. task interaction *F*_(1,18)_ = 5.357, *P* = 0.0327. Multiple comparisons between social reward vs. neutral context *P* = 0.0003, social reward context vs. preferred context; *P* = 0.0004, *n* = 10 neurons out of a total of 37 recorded neurons, *n* = 5 mice.

### The vH represents emotional contexts in a valence-specific manner

Next, we asked whether vH contextual representations may differ according to valence. To address this question, we subjected mice to a discriminative contextual fear conditioning procedure involving two contexts: context A in which mice received electrical foot shocks and context B where no foot shocks were delivered (*n* = 24 mice, Fig. 4A). Following contextual fear conditioning, mice exhibited high freezing levels in context A at the memory test session whereas freezing levels in context B remained low, indicating that mice formed a selective contextual fear memory (Fig. 4B).

**Figure 4.**
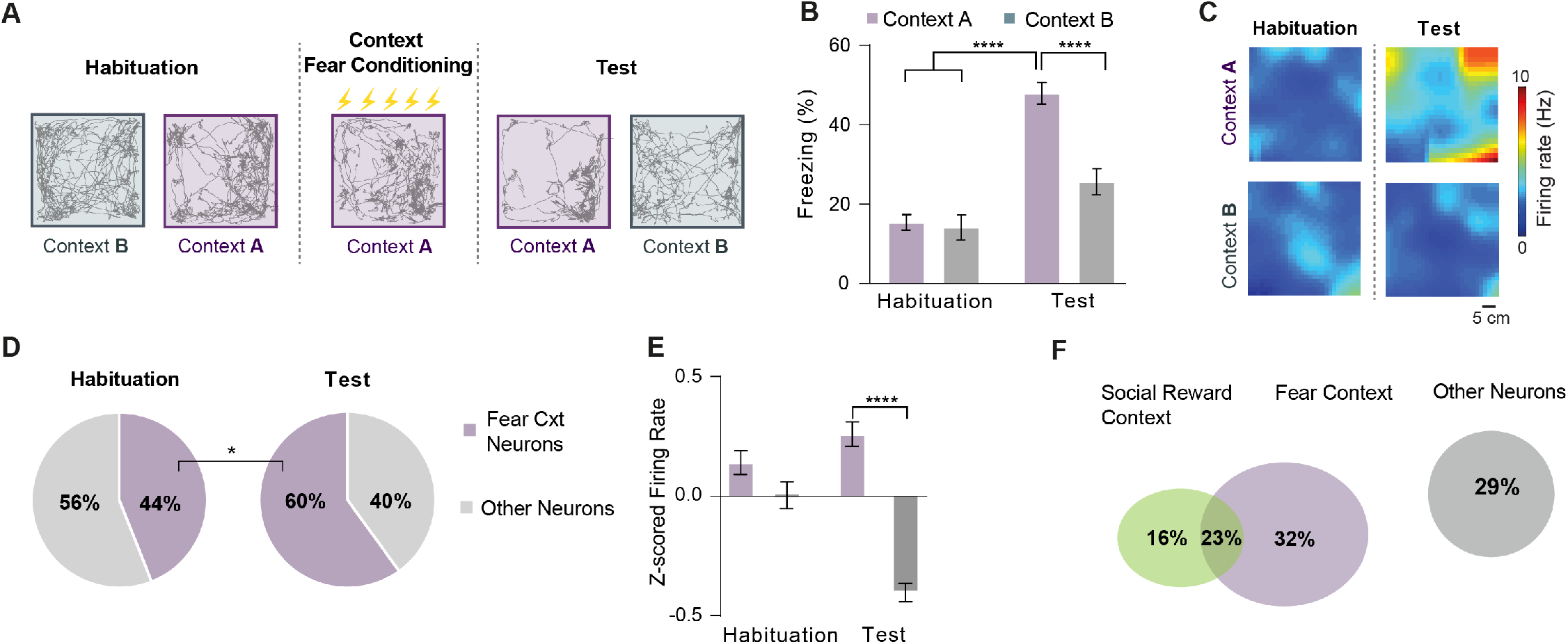
The vH represents emotional contexts in a valence-specific manner. **(A)** Behavioural protocol and representative mouse trajectories during the habituation, fear conditioning and test sessions for the contextual fear memory task. **(B)** Freezing levels during habituation and test sessions. Two-way repeated measures ANOVA between groups, context x session factor *F_(1,46)_* = 14.98, *P* = 0.0003. Multiple comparisons between context A and context B during test *P* < 0.0001 and habituation and test at context A *P* < 0.0001, *n* = 24 mice. **(C)** Example of spatial firing rate maps of a fear context neuron at the habituation and test sessions in context A and B. **(D)** Comparison of fear context neurons proportions at habituation and test sessions. Chi-square test between groups, *P* = 0.0235, *n* = 24 mice. **(E)** Averaged normalized firing rates of fear context neurons during the habituation and test sessions in context A and B. Two-way repeated measures ANOVA between groups, session x context effect *F*_(1,226)_ = 18.50, *P* < 0.0001. Multiple comparisons between Context A vs. Context B at the Test session *P* < 0.0001, *n* = 114 neurons, *n* = 24 mice. **(F)** Overlap of neurons with preferential activity in the social reward context and fear context, *n* = 154 neurons in total, *n* = 23 mice.

An examination of the contextual representations at the memory test session revealed that the majority of vH neurons were selectively active in context A (corresponding to fear context neurons) concomitant with an increase in the fraction of vH neurons representing context A compared to the habituation session (Fig. 4C, D). During the memory test session, fear context neurons showed increased firing selectively in context A compared with context B, which was not observed during habituation (Fig. 4E). This finding suggests that, similar to our findings with social reward contextual conditioning, the vH preferentially represents the emotional context after learning. Of note, the activity of vH neurons was not correlated with the speed of movement during contextual fear conditioning (Supplementary Fig. 3A). Moreover, using a SVM to decode context identity in individual mice, we found that the classification accuracy was higher for the model trained on correct compared to randomized labels suggesting context was also represented at the population level (Supplementary Fig. 3B). To determine if the presence of an aversive reinforcer was necessary for the formation of context-discriminating activity and memory in the vH, a control paradigm using a neutral stimulus (i.e. two foot shocks of 0.06 mA) was conducted in a separate cohort of mice (n = 3 mice, Supplementary Fig. 4A). Mice did not exhibit a contextual fear memory during this control conditioning (Supplementary Fig. 4B), but when subjected to the classical contextual fear conditioning, they expressed contextual fear memory associated with the formation of fear context neurons (Supplementary Fig. 4 C and D). This indicates that selective contextual representations are formed in the vH during contextual fear conditioning.

Finally, we examined the extent to which context-discriminating vH neurons overlapped in the social reward and fear paradigms. We found that a rather small proportion of vH neurons (23%) exhibited social reward and fear context activity in both paradigms, while a larger proportion of neurons were selectively activated in the social reward or fear contexts (48%). This suggests that the vH can discriminate contextual valence by two partly non-overlapping neuronal populations (Fig. 4F).

### Social reward contextual memory involves locus coeruleus projections to the vH

Next, we addressed the potential neuromodulatory inputs to the vH that may play a role in regulating contextual associations. Since the LC has been implicated in facilitating spatial learning (Kempadoo et al., 2016; Takeuchi et al., 2016) and supporting the enrichment of place cells near reward zones (Kaufman et al., 2020), we investigated whether LC afferents to the vH may be involved in social reward contextual conditioning. The LC of tyrosine hydroxylase (TH)-Cre mice were bilaterally injected with a Cre-dependent adeno-associated virus (AAV) carrying the inhibitory opsin archaerhodopsin (eArch3.0) fused with yellow fluorescent protein (EYFP), followed by bilateral optic fibre implantion into the vH (Fig. 5A and Supplementary Fig. 5A). Four weeks later, mice underwent the social reward CPP paradigm to evaluate the necessity of LC terminal activity in the vH for the formation of a social reward contextual memory. During training sessions, light was delivered upon the entry of mice into the social reward context in a closed-loop manner (Fig. 5B). Optogenetic suppression of LC activity in the vH did not affect the time spent in the social reward context or the social reward zone during training, as indicated by similar discrimination scores between control and Arch mice (Fig. 5C and Supplementary Fig. 5B). However, suppressing LC activity in the vH during training resulted in a significant decrease in the preference for the social reward context at the Post-Train Test session, suggesting an impairment in social reward contextual memory (Fig. 5C and Supplementary Fig. 5B). These results point to the necessity of LC projections to the vH in associating social reward stimuli with contextual information to form a social reward contextual memory. To test for a valence-specific effect, we evaluated the impact of optogenetically suppressing LC terminal activity in the vH during contextual fear conditioning in mice previously tested in the social reward CPP paradigm, but we did not observe significant changes in contextual fear memory (Supplementary Fig. 6A,B).

**Figure 5.**
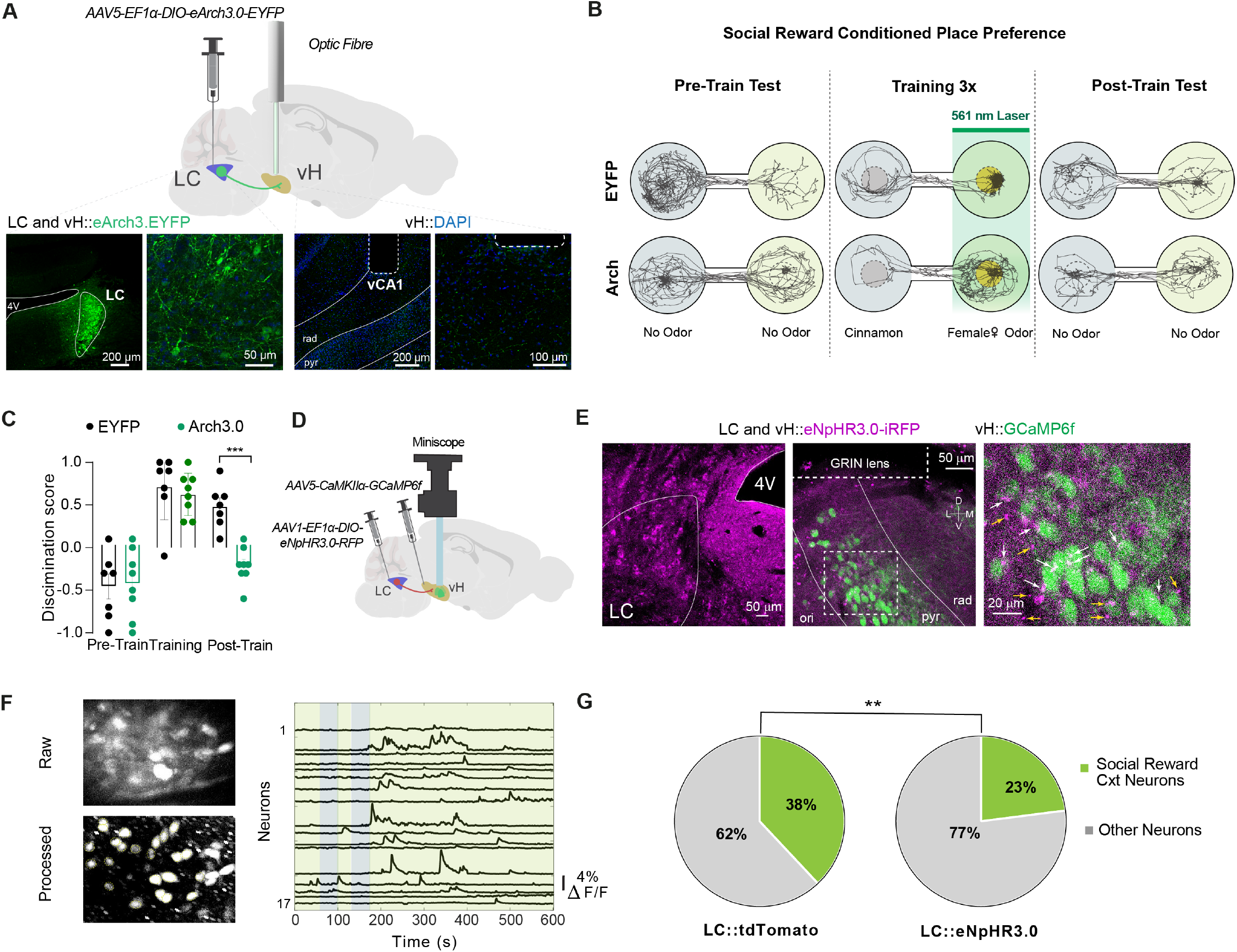
Social reward contextual memory involves locus coeruleus projections to the vH. **(A)** Schema of the experimental strategy (top) with representative confocal microscopy images (bottom) of LC somatic labelling (left) and LC projections labelling in the vH (right). **(B)** Schema of the optogenetic inhibition protocol during the social reward CPP. **(C)** Discrimination score during the social reward CPP. Two-way repeated measures ANOVA between groups, Condition x session effect *F*_(2,24)_ = 4.254, *P* = 0.0263. Multiple comparisons between groups EYFP vs. eArch3.0 during Post-Train Test session, *P* = 0.0002, *n* = 7 EYFP, *n* = 8 eArch3.0 mice. **(D)** Schema of the experimental strategy using miniscopes for *in vivo* calcium imaging of vH pyramidal neurons during social reward CPP. **(E)** Representative confocal microscopy images of LC somatic labelling (left) and LC projections labelling in the vH with GCaMP6f expression in pyramidal neurons at low (middle) and high (right) magnification. Yellow arrows indicate LC varicosities and white arrows indicate LC varicosities superimposed on GCaMP6f positive pyramidal neurons. **(F)** Left, raw and processed images of miniscope-acquired calcium signals in vH pyramidal neurons. Right, sampling calcium traces of pyramidal neurons during the Post-Train Test session of the social reward CPP. **(G)** Comparison of proportions of social reward context neurons of tdTomato and eNpHR3.0 mice during the Post-Train Test session. Chi-square test between groups, *P* = 0.0096, tdTomato: *n* = 84 neurons, 2 mice, eNpHR3.0: n = 179 neurons, 2 mice.

We hypothesized that LC activity in the vH had an impact on the development of context-discriminating activity in the social reward CPP paradigm. To test this hypothesis, we used head-mounted miniscopes to perform calcium imaging of vH pyramidal neurons together with optogenetic inhibition of LC terminals in the vH while performing the social reward CPP paradigm (Cai et al., 2016; Ghosh et al., 2011)(Fig. 5D,F). To do so, TH-Cre mice were injected with a conditional AAV carrying the inhibitory opsin eNpHR3.0 into the LC, combined with a second unilateral AAV injection to express the calcium indicator GCaMP6f in vH pyramidal neurons using a CaMKII promoter. Next, a gradient refractive index (GRIN) lens was unilaterally implanted above the injection site in the vH pyramidal layer (Fig. 5D, E and Supplementary Fig. 5C). LC neurons projecting to the vH were inhibited as mice entered and occupied the social reward context during the social reward CPP training sessions. We then analysed the proportion of social reward context neurons in the vH during the Post-Train Test session and found that mice with inhibition of LC projections in the vH showed a significant reduction in social reward context neurons during the Post-Train Test compared with controls (Fig. 5G). Collectively, these results support the idea that the LC modulates social reward contextual memories by promoting the formation of social reward context neurons in the vH.

## Discussion

We found that the vH supports the formation of contextual memory for social rewards by generating context-specific spatial representations. These representations emerged upon learning, involved a preferential remapping of place fields to the social reward context, and depended on the presence and valence of reinforcers. A circuit-based strategy identified LC projections to the vH as a key modulator of social reward contextual representations and memory.

Animals must be able to detect and discriminate stimuli according to their valence to support adaptive behaviour and memory processes. In various species, chemosensory cues from females are considered naturally rewarding and highly attractive to males, resulting in the activation of the dopaminergic reward system (Malkesman et al., 2010). In our study, sexually naïve males were exposed to female odorant/pheromonal stimuli through the soiled bedding of unfamiliar females in a specific context during a social reward contextual conditioning procedure. While a role of the ventral CA1 and dorsal CA2 regions of the hippocampus has been highlighted in the storage of social memories related to conspecifics (Hitti and Siegelbaum, 2014; Meira et al., 2018; Okuyama et al., 2016; Phillips et al., 2019; Sun et al., 2020), much less is known about whether selective contextual representations in the vH are formed for social reward and the underlying neuronal dynamics and circuits.

We discovered that a selective population of vH neurons, i.e. the social reward context neurons, selectively tune their activity to the social reward context upon learning, consistent with the fact that the hippocampus (both its dorsal and ventral subdivisions) displays context-specific activity patterns after reward conditioning (Ciocchi et al., 2015; Komorowski et al., 2013; Sjulson et al., 2018; Trouche et al., 2016). Social reward contextual learning produced a remapping of the spatial activity fields of vH neurons, in agreement with former studies showing that vH neurons change their spatial firing activity in the presence of unconditioned olfactory cues, such as predator odour (Keinath et al., 2014), as well as odours associated with reward (Komorowski et al., 2013). Specifically, in a biconditional odour discrimination task involving sucrose rewards in two distinct contexts, vH CA3 neurons were found to be originally active across both contexts but became context-selective as animals learned the odour-reward contingencies (Komorowski et al., 2013). This corroborates our finding of a large proportion of vH neurons exhibiting place fields covering both contexts before conditioning, but that remapped towards the social reward context after training.

Early observations that information is more readily recalled in the environment where learning had occurred led to classical psychological theories on the ability of spatial contexts to regulate memory retrieval (Godden, 1975). Strikingly, in our experiments, the fraction of social reward context neurons correlated with contextual memory retrieval, suggesting that hippocampal contextual representations imbued with positive value influence the strength of memory retrieval. Of note, we observed that a small proportion of vH social-context neurons were further activated in the social reward zone. This suggests, on one hand, that soiled bedding containing female odours and pheromones is likely perceived as a social reward by males and, on the other hand, that the vH is predominantly involved in encoding spatial representations related to contextual memories. Nevertheless, the social reward responsiveness of vH neurons indicates that converging information from odour-driven long-range synaptic inputs to the vH, such as from the piriform cortex (Poo et al., 2022), as well as other inputs from brain regions coding social salience, such as the amygdala (Felix-Ortiz and Tye, 2014), and dCA2 (Meira et al., 2018) could instruct vH contextual representations during learning.

Contextual representations in the hippocampus are thought to be transmitted to downstream regions where they are integrated with stimulus identity, associated values, and behavioural outcomes (Maren et al., 2013). Our lab and others have found spatial and valence coding in the ventral hippocampus (Ciocchi et al., 2015; Forro et al., 2022; Royer et al., 2010), hinting at the possible confluence of the animal’s current location in space and its emotional state. Strikingly, vH neurons exhibiting context-dependent representations following learning poorly overlapped in paradigms of opposite valence, in line with previous studies showing functional heterogeneity within vH pyramidal neurons and their projection targets (Bienkowski et al., 2018; Gergues et al., 2020; Shpokayte et al., 2022). We predict that these contextual- and valence-specific neuronal representations in the vH likely diverge in their long-range synaptic targets. Consistent with this prediction and to support context-mediated emotional memory selection, social reward context neurons in the vH may preferentially project to the NAc (Ciocchi et al., 2015; Okuyama et al., 2016; LeGates et al., 2018; Zhou et al., 2019) or medial prefrontal cortex (Phillips et al., 2019; Sun et al., 2020) to signal that the context is rewarding in nature, while fear context neurons may preferentially target the basolateral amygdala (Jimenez et al., 2018; Xu et al., 2016) to signal that the context is aversive. Thus, selective contextual representations of positive and negative emotions in the vH may gate or even facilitate adaptive behavioural responses mediated by downstream structures.

It is widely established that the LC is the primary source of noradrenaline in the brain, with several studies showing the impact of the LC noradrenaline system on cognitive and behavioural processes, such as the regulation of attention and working memory, goal-directed and motivated behaviour, and emotional memories(Poe et al., 2020; Sara and Bouret, 2012). There is also evidence that dopamine released by LC neurons in the dorsal hippocampus is critical for memory formation (Chowdhury et al., 2022; Kaufman et al., 2020; Kempadoo et al., 2016; Wagatsuma et al., 2018). Additionally, former studies showed that dopamine plays a role in the manifestation of male sexual behaviour in terms of motivation and copulatory ability (Beny-Shefer et al., 2017; Dai et al., 2022). In our study, suppressing LC activity in the vH impaired the formation of social reward contextual representations and memory without affecting motivation to approach the social reward zone or context during learning. Instead, LC-vH inhibition may have disrupted the association of the social reward stimulus with the animal’s contextual location.

The release of noradrenaline or dopamine from the LC can have several implications for the function of vH circuits and the formation of emotional memories. One possibility is that these neuromodulators fulfil a pure gain-control function to signal stimulus intensity thereby regulating learning strength. Successful learning requires plasticity mechanisms at the level of single neurons and synapses, which are thought to lead to the stabilization of place fields to mediate spatial memory formation (Bliss and Lomo, 1973; Buzsáki, 2015; Martin et al., 2000; Kentros et al., 1998; McHugh et al., 1996). The hippocampus contains dopamine and adrenergic receptors whose activation by their corresponding neurotransmitters promotes changes in synaptic efficacy, thus contributing to memory formation and consolidation (Broussard et al., 2016; Hu et al., 2007; Kwon et al., 2008; Liu et al., 2017; Rocchetti et al., 2015; Wang et al., 2010; Kentros et al., 1998; Li et al., 2003; McNamara et al., 2014; Murchison et al., 2004). The role of noradrenaline in learning is further supported by results showing that the stimulation of LC projections to the dorsal hippocampus promotes spatial learning (Kempadoo et al., 2016; Takeuchi et al., 2016) and in the case of spatial reward learning, prompts place cell enrichment adjacent to the reward location (Kaufman et al., 2020). Altogether, and in line with the literature, our study suggests that LC transmission to the vH promotes the formation of stable spatial representations in the social reward contextual conditioning paradigm. Yet, future studies are needed to evaluate which neuromodulators released by the LC are involved in contextualization of a social reward memory and by which mechanisms.

Finally, we found that LC projections to the vH were necessary for associating contexts with a social reward but not with an aversive stimulus. These results were rather unexpected as there is evidence of the involvement of the LC in fear memory (Giustino et al., 2019; Uematsu et al., 2017; Wagatsuma et al., 2018). For instance, during auditory fear conditioning, optogenetic inhibition of the LC coinciding with shock delivery impaired fear learning (Uematsu et al., 2017), while pharmacogenetic activation of noradrenergic neurons in the LC has been found to induce fear relapse (Giustino et al., 2019). A caveat to consider in our experiments is that the social reward CPP and contextual fear conditioning procedures differ in nature. In the social reward CPP paradigm, mice were exposed to both contexts simultaneously and could freely choose which context to explore. In contrast, in the contextual fear conditioning paradigm, the contextual exposures to Context A and B were separated in time and space. This may suggest that LC projections to the vH may be crucial for behaviours involving an approach-avoidance conflict as in the social reward CPP paradigm. Another caveat is that the foot shock reinforcer may be more strongly perceived than the social reward reinforcer, leading to the formation of more fear-context neurons in the vH, which would be sufficient for intact learning. Consistent with this idea, our contextual fear conditioning paradigm led to the formation of more fear context neurons (roughly 60%) compared to social reward context neurons (roughly 40%) in the social reward CPP paradigm.

In conclusion, our observations together with prior studies identify the vH as a crucial node for contextual emotional memories as part of a wide-ranging neuronal circuitry integrating converging spatial and emotional information. Our study, in mice, provides insight into how spatial contextual representations in the hippocampus are imbued with emotional values to ensure context-dependent memory selection. These findings may have implications for neuropsychiatric disorders like substance abuse or post-traumatic stress disorder, where the improper linking of emotional stimuli to the environment leads to maladaptive behaviours that are excessive or indiscriminate (Liberzon and Abelson, 2016).

## Material and Methods

### Mice

Male C57BL/6J mice (Janvier Labs, France) or tyrosine hydroxylase Cre mice (Jackson Laboratories, B6.Cg-7630403G23RikTg(TH-Cre)1Tmd/J) aged 3-4 months old were used for experiments. Mice were grouped housed (2-4 mice per cage) in a 12h light/dark cycle and provided with food and water ad libitum. Mice used for electrophysiology experiments were individually housed beginning three days before surgery. All behavioural experiments were conducted during the light cycle. All animal procedures were executed following institutional guidelines and were approved by the prescribed authorities (Veterinary Office of the Canton of Bern, Switzerland).

### Surgery

Mice were anaesthetized with isoflurane (induction 3%, maintenance 1.5%) in oxygen at a flow rate of 1 L/min throughout the procedure. Core body temperature was kept at 37 °C by a feed-back controlled heating pad (Harvard Apparatus GmbH, Breisgau, Germany). Ophthalmic cream was applied to avoid eye drying. The mice were positioned into a stereotaxic frame (David Kopf Instruments, California, USA), and local analgesia was applied by injecting a mixture of 2% of lidocaine (Streuli Pharma, Uznach, Switzerland) and 0.5% bupivacaine (Aspen Pharma Schweiz, Baar, Switzerland) subcutaneously under the scalp. Anesthesia was confirmed by detecting deep breathing, slow heart rate, and lack of the toe pinch reflex. The skull was uncovered after a skin incision, and Bregma and Lambda were aligned. A craniotomy was made on the skull above the region of interest. The dura matter was removed, and saline solution and haemostatic porcine gelatin sponges (Equimedical) were applied to the brain surface to avoid edemas.

#### In vivo electrophysiology

C57BL6/J mice were unilaterally implanted in the vH with a custom-made microdrive (Axona Ltd, St Albans, UK) holding eight tetrodes targeted at the following coordinates: AP 3.0 mm, ML ± 3.1 mm, DV 4-4.3 mm from the bregma. The tetrodes consisted of eight individually insulated, gold-plated tungsten wires (12.7 µm inner diameter, impedance 100-300 kΩ, California Fine Wire Company, California, USA) and were placed in a stainless steel guide cannulas and attached to a 32-pin connector (Omnetics Connector Corporation, Minnesota, United States). Two stainless steel screws implanted above the cerebellum were used as ground electrodes. Next, the exposed parts of the tetrodes were protected using sterile wax. Then the microdrive was fixed to the skull with light-cured dental adhesive (Kerr, OptiBond Universal, Kloten, Switzerland) and dental cement (Ivoclar, Tetric EvoFlow, Schaan, Principality of Liechtenstein). Mice were given Carprofen (5 mg/kg, subcutaneous) for three days after surgery.

#### Optogenetics experiments

The LC (AP -5.45 mm, ML ± 0.9 mm, DV -3.65mm from Bregma) of TH-Cre mice were injected bilaterally into the with AAV2/5-EF1α-DIO-eArch3.0-EYFP or AAV2/5-EF1α-DIO-EYFP (both viruses from UNC vector core, 0.3 µl of viral solution) via a glass micropipette (Blaubrand, Brand, Wertheim, Germany) and a microinjector Picospritzer III (Parker Hannifin, Ohio, USA). The micropipette was left in place for 10 minutes after infusion to avoid backflow. Optic fibres were lowered into vH at coordinates of AP 2.9 mm, ML 3.3 mm and DV -4.0 mm at at low speed of 1mm/min. Optic fibres were constructed by gluing a piece of multimodal optical fibre (200 µm, 0.37 NA, Thorlabs, New Jersey, USA) to a 1.25 mm diameter, 230 µm bore ceramic ferrule (Senko, Pennsylvania, USA). The optic fibre implants were secured to the skull with stainless steel screws and light-cured dental adhesive and dental cement. Mice were allowed to recover for at least four weeks before behavioural assays to ensure sufficient expression of the viral construct.

#### Calcium imaging

Mice were unilaterally injected in the LC (AP -5.45 mm, ML ± 0.9 mm, DV -3.65 mm from Bregma) with AAV1-EF1α-DIO-eNpHR3.0-iRFP (University of Zurich viral vector facility) or AAV8-EF1α-Flex-tdTomato (UNC vector core) using a glass micropipette (0.3 µl of viral solution). The micropipette was left in place for 10 minutes after infusion and then slowly retracted to avoid backflow. Then, mice were injected with 0.5 µl of AAV2/5-CaMKIIα-GCaMP6f (Addgene, #100834) into the vH (AP -3.28 mm, ML +3.45 mm, DV -4.0 mm from Bregma). The micropipette was left in place for 45 minutes after infusion. Three skull screws were inserted around the implantation site. The GRIN lens (0.6 mm diameter) was slowly lowered to DV -4.0 mm by using a leading 21 gauge needle attached to a custom-made stereotaxic guide that allowed precise placement of the lens. The lens was fixed to the skull surface with light-cured dental adhesive and dental cement. The dorsal surface of the skull and the three bone screws were cemented with the GRIN lens to ensure the implant’s stability.

Three weeks later, mice with suitable virus expression were fixed in a stereotaxic frame under anaesthesia to attach an aluminium baseplate for the miniscope above the GRIN lens. After finding the best field of view, the baseplate was cemented to the skull and a cap was used to protect the GRIN lens from dust. Mice wore a dummy miniscope for 1 week to adapt to the additional weight on the head before behavioural tests and calcium signal recordings.

### Behaviour

Behavioural tests were conducted under room lighting (100 lux) and recorded with an overhead webcam video camera at 30 Hz. Tracking and scoring were done using ANY-maze (Stoelting, IL, USA). Between behavioural sessions, mice stayed in their home-cages in a sound-attenuating chamber.

#### Social place preference conditioning

This paradigm was implemented in a two-chamber apparatus made of dark grey plastic and consisting of two circular contexts, each measuring 25 cm in diameter, joined together with a 10 cm narrow middle chamber. Each context had either black and white vertical stripes or all-white wall patterns and different textured floors. The social conditioning place preference test protocol was adapted from (Beny-Shefer et al., 2017) and consisted of three stages conducted over two days: on day 1, (1) a 10-minute habituation session in which an 8.5 cm diameter petri dish containing clean woodchip bedding was placed in the centre of each context to evaluate side preferences; (2) two hours later, three 10 minutes learning sessions separated by 1 hour, in which female-bedding was presented in the non-preferred context (social context) and clean bedding mixed with a neutral odour (1% of cinnamon, (Aqrabawi and Kim, 2018) in the other context (non-social context); and on day 2, (3) a 10-minute memory test session in which clean bedding was placed in both contexts. The apparatus was cleaned with 70% ethanol solution and dried between sessions. The time spent in each context was used to calculate a discrimination score as follows: time in social context minus time in non-social context divided by their sum. Positive values showed that the animal exhibits a preference to the social context. 23 mice were used in the social place preference conditioning paradigm (compared to 24 mice in contextual fear conditioning) because one animal exhibited poor exploration due to cable twisting and was thus removed from the analysis.

A subset of mice were exposed to a control ‘conditioning’ procedure prior to the social place preference conditioning protocol. This control ‘conditioning’ paradigm used a two-chamber apparatus of the same size as previously described, however, with squared shapes with different visual patterns on walls and textured floors. Experiments were performed in a different position in the room exposing mice to different landmarks. The neutral ‘conditioning’ protocol used clean bedding mixed with 1% of rosemary in one context and clean bedding mixed with 1% of saffron in the other context during the three learning sessions. The rest of the protocol was identical to the social place preference conditioning.

#### Contextual fear conditioning

This behavioural paradigm took place in two different squared contexts of transparent Plexiglas (26 cm x 26 cm). These two contexts displayed different floor textures (grid vs. smooth floor), wall patterns, distal cues, and odours (70% ethanol or 1% acetic acid). The fear context contained a grid floor made of stainless steel rods through which scrambled electrical shocks were delivered by a shock controller (Coulbourn Instruments). Foot-shock intensity was adjusted to 0.5 mA before each experiment. The two arenas were cleaned before and after each session with 70% ethanol or 1% acetic acid, respectively.

Mice were first habituated to the two contexts (A and B) in 10 minute sessions separated by a 1 hour interval. Two hours later, mice were placed in the fear conditioning context (A) and received five electrical foot-shocks (0.5 mA, 1 s duration) delivered through the floor grid. Four hours later, mice were tested for contextual fear memory by returning them to the fear conditioning context (A) for 10 minutes. Two hours later, mice were placed into the neutral context (B) to assess the animal’s ability to discriminate the contexts. Freezing levels were automatically detected by ANY-maze software which tracked body positions at 30 Hz.

A subset of mice was exposed to a control ‘conditioning’ procedure followed by the classical contextual fear conditioning paradigm. In the control ‘conditioning’ paradigm, the contexts had equivalent dimensions with one context being circular in shape and differed in their visual cues and floor texture. In this control conditioning paradigm, mice received two weak foot-shocks of 0.06 mA. The rest of the protocol was identical to the contextual fear conditioning protocol. Following control ‘conditioning’, the mice were subjected to the classical contextual fear conditioning paradigm as previously described.

### Single-unit electrophysiological recordings in freely-moving mice

Single-unit electrophysiological recordings were perfomed on chronically implanted mice. After recovery from the microdrive implantation surgery, each animal was familiarized with the testing room and handling procedures which involved cable tethering to the recording system. Tetrodes were progressively lowered to the vH pyramidal layer over several days by using sharp-wave ripples and theta oscillations as electrophysiological hallmarks. Twenty-four hours later, mice underwent behavioural testing as defined above. Electrophysiology and behavioural tracking data were synchronized using TTL signals sent by Anymaze to the recording system to timestamp single-camera frames or shock delivery –in the case of the contextual fear conditioning procedure.

The electrophysiological data were acquired using an Intan RHD2000 Evaluation Board and 32 channel headstage at a sampling rate of 20 kHz. The signal was amplified and a band-pass filter (500-5000 Hz) was applied offline to extract the spikes. Spikes were identified using a threshold of 5 standard deviations above the root mean square signal (0.2 ms sliding window) of the filtered signal. 32 data points (1.6 ms) were sampled for each spike waveform.

Principal component analysis (PCA) was applied to the waveforms to extract the first three components per tetrode channel. Identified spikes were automatically sorted using KlustaKwik (http://klustakwik.sourceforge.net/) followed by manual adjustment of the clusters using the software Klusters (http://neurosuite.sourceforge.net/) to acquire well-isolated single-units based on temporal cross-correlations, spike waveforms and refractory periods. The stability of single-units was confirmed by inspecting spike features throughout the recording sessions.

### Calcium imaging

Imaging sessions in freely-moving mice began five days after baseplating. Mice were briefly anaesthetized (< 2 min) to attach the miniscope to the baseplate for each imaging day. Mice were allowed to recover from the brief anaesthesia for 60 minutes before the behavioural protocol started. Calcium imaging was performed using a custom-made dual-color miniscope capable of simultaneous calcium imaging and optogenetic manipulation (Srinivasan et al., 2019). The power of the blue laser used for GCaMP6f excitation (488 nm, Cobolt, Solna, Sweden) was set to 1 mW at the level of the miniscope objective. The power of the red-orange laser used for eNpHR3.0 excitation (594 nm, Cobolt) was set to 8-10 mW at the level of the miniscope objective. The 488 nm laser was triggered by a TTL signal from ANY-maze at the beginning of each recording session. The 594 nm laser was switched on only when the mouse entered the social reward context. Calcium imaging videos were recorded at 20 Hz as uncompressed .avi files (1000 frames per file) by using a data acquisition (DAQ) box which is triggered by an external TTL pulse from ANY-maze to allow for simultaneous acquisition of calcium imaging and behavioural videos. The excitation power for GCaMP6f was determined in prior tests based on the most optimal signal to noise ratio and was maintained throughout all the imaging/behavioural sessions.

### Optogenetic stimulation

To optogenetically manipulate LC projections to the vH (starting four weeks after viral injections), a laser (Cobolt, Solna, Sweden) generating green light (561 nm) was attached to an optical rotary joint (Doric Lenses, Québec, Canada) to support the unrestricted movement of mice during the behavioural tests. The optical rotary joint was connected to a light splitter (Doric lenses, Québec, Canada) to allow bilateral light delivery to two patch cables (Doric Lenses, Québec, Canada) which were in turn connected to the implanted optic fibres through a ferrule-sleeve system (Senko, Pennsylvania, USA). Light illumination was automatically controlled by ANY-maze based on the animal’s body position in the behavioural apparatus. Laser power was measured with a power-meter (Thorlabs, New Jersey, USA) before each behavioural sessions to ensure a power of 10-15 mW at the implanted fibre tip. Before the beginning of the behavioural paradigm, mice were first connected to the patch cables for 10 minutes for habituation.

### Histology

For electrophysiological experiments, mice were anaesthetized with isoflurane and tetrode positions were marked by electrolytic lesions (30 µA for 10 seconds). Then, mice were anaesthetized using an intraperitoneal injection of a ketamine/xylazine mixture (5% ketamine, Vetoquinol, Magny-Vernois, France; 2.5 % xylazine, Streuli Pharma, Uznach, Switzerland). Mice were then transcardially perfused with ice-cold phosphate-buffered saline (PBS, Roche, Basel, Switzerland) followed by 4% paraformaldehyde (PFA, Carl Roth, Karlsruhe, Germany). Implanted tetrodes or optic fibres were kept in the brain for at least 24h of post-perfusion fixation in 4% PFA at 4 °C before being detached from the brain. Subsequently, brains were removed from the skull and post-fixed for 24h at 4 °C. Brains were sliced (50 µm thick coronal sections) using a vibratome (VT1000 S, Leica Microsystems, Wetzlar, Germany). Sections from electrophysiological or calcium imaging experiments containing the vH were examined with a fluorescent microscope (M205 FCA, Leica Microsystems, Wetzlar, Germany) using a 2.0x magnification objective. The location of the tetrodes or the GRIN lens was manually verified by comparing the location to the mouse atlas (Franklin and Paxinos 2017). If the tip of the tetrodes or the GRIN lens was not located in the vH, the animal was excluded from further analysis. Afterwards, sections were stored in PBS containing 0.05% sodium azide.

#### Immunohistochemistry

Free-floating sections were washed in PBS and blocked with 5% normal donkey serum (ab7475, Abcam, Cambridge, United Kingdom) in PBS containing 0.1% Triton X-100 (PBS-T) at room temperature for 2 hours. Afterwards, sections were incubated with rabbit anti-GFP polyclonal antibody (1:1000, ab6556, Abcam, Cambridge, United Kingdom) or rabbit anti-RFP polyclonal antibody (1:1000, 600-401-379, ThermoFisher, Massachusetts, USA) and chicken polyclonal anti-GFP (1:1000, ab13970, Abcam, Cambridge, United Kingdom,) in PBS-T for 48 hours at 4 °C followed by three washings in PBS. Then, the slices were incubated with Alexa Fluor 488 conjugated donkey anti-rabbit secondary antibody (1:1000, A32790, Invitrogen, Massachusetts, USA) or donkey anti-chicken AlexaFluor-488 (1:1000, 703545145, Jackson ImmunoResearch Laboratories, Cambridge, UK) and donkey anti-rabbit AlexaFluor-564 (1:1000, A21207, Invitrogen, Massachusetts, USA) for 2 hours at room temperature. Finally, the slices were washed three times in PBS and incubated in DAPI solution (MBD0015, Sigma-Aldrich, Missouri, USA) to label cell nuclei, subsequently washed another three times with PBS, mounted onto microscope slides with aqueous mounting medium (Aqua-Poly/Mount, Polysciences, Pennsylvania, USA) and stored at 4 °C. Immunolabelled sections were imaged with a 20x objective on a slide scanning microscope (3DHistech, Pannoramic 250 Flash II, Budapest, Hungary) or using a confocal microscope (LSM 880, Zeiss, Jena, Germany). Brain structures were defined according to Franklin and Paxinos (Mouse brain atlas 2017) and the location of viral transduction and optic fibre implantations was verified.

### Calcium signal processing

Calcium imaging videos were analysed using a custom Matlab code (Grewe et al., 2017). Videos from multiple sessions were concatenated and downsampled by a binning factor of 4 resulting in a frame rate of 5Hz, and lateral brain movement was motion-corrected using the Turboreg algorithm (Thévenaz et al., 1998). Fluorescent traces were extracted by applying automatically detected individual cell filters based on combined principal and individual component analysis (PCA/ICA) as described in (Mukamel et al., (2009). To control for non-inclusion of split neurons in our analyses, we identified pairs of neurons with highly correlated activity (Pearson correlation > 0.7) and are spatially close (centroid distance < 20 pixels) and excluded one of the neurons for each pair. Identified putative neurons were then sorted via visual inspection to select neurons with appropriate somatic morphology and Ca^2+^ dynamics consistent with signals from individual neurons.

### Data Analysis

#### Categorization of social reward context and fear context neurons

We used a bootstrap analysis to define whether individual vH neurons were significantly active in the conditioned emotional context at memory test to categorize social reward context neurons or fear context neurons. First, for each well-isolated neuron, a spike firing-by-time vector was constructed by binning its spikes in 20 ms bins and then smoothed by convolution with a Gaussian kernel (σ = 20 ms). Then, a Student’s t-statistic value was calculated based on the difference in firing rate activity between the social reward context and its corresponding neutral context (or fear context and its corresponding neutral context in the case of fear conditioning). This calculation was then repeated 100000 times by resampling the data with replacement which led to a bootstrap distribution of resampled data. The original Student’s t-statistic value was compared to the bootstrapped distribution. Neurons whose observed statistical value fell at the extremes of the surrogate distribution (*P* < 0.001) were considered as significantly responsive, with a positive value showing higher activity in the neutral context and a negative value showing higher activity in the social context or respectively in the fear context. To detect neurons excited in the social reward zone (i.e. petri dish zone containing female bedding), we compared firing rate activity inside and outside of the social reward zone or neutral context during training using the same bootstrap method.

#### Place maps

For each neuron, place maps were generated based on the mean firing rate in each spatial bin. Only timepoints with an instantaneous velocity greater than 2 cm/s measured from the body’s centre point were included for analysis. First, each spike was assigned to a 1 x 1 cm maze spatial bin. The firing rate was then calculated by dividing the number of spikes occurring in each bin with the occupancy time in that bin. The place maps were then smoothed by convolving them in the X and Y dimension with a Gaussian kernel (σ = 3 cm). For place field remapping, the place field localization was evaluated using the location of the peak firing rate within the place field. Place field stability was measured by performing pixel-by-pixel Pearson’s r correlations between the Pre-Train and Post-Train sessions.

#### Contextual firing rate analysis

For each neuron, the firing rate was first calculated by binning spikes in 20 ms bins and dividing by the time interval. For analysis of activity time-locked to context entries, the firing rates of individual neurons were further binned into 2 s intervals, then Z-scored using the mean and standard deviation of the baseline firing rate prior to context entry. To calculate the normalized mean firing rate in each behavioural paradigm, a z-score was computed using the mean and standard deviation in the mean firing rates across all sessions.

#### Context decoding

Smoothed firing rates in 20 ms bins were obtained for each neuron in each context. For individual neurons, the firing rate was shuffled and a random sample of 60 seconds was selected for each context and labelled according with the context identity. The data were then randomly split (30/70) into training and hold-out test samples with a balanced number of labels. Training samples were used to train a binary linear classifier (support vector machine [SVM], Matlab Classification Learner application) with cross-validation (10-fold) and then applied to test samples with accuracy calculated as the percentage of correctly labelled samples. For the SVM calculated for individual mice, only mice with more than 10 recorded neurons were incorporated in the analysis.

### Statistics

Analyses were performed using custom scripts written in MATLAB (MathWorks). Statistical analyses were carried out using Prism 9 (Graphpad software). All datasets were tested for normality using the Kolmogorov-Smirnov test. All null-hypothesis tests were two-tailed. ANOVAs were followed by post hoc tests if a main effect was observed. Post hoc multiple comparisons were done using the Bonferroni correction. Chi-square tests were conducted to compare differences in proportions between groups. P-values for statistical significance are reported in the figure legends. Box and whisker plots show median, interquartile range (25th and 75th percentiles). All data are shown as mean ± S.E.M. Asterisks in the figures represent *P*-values with the following thresholds: * *P* < 0.05; ** *P* < 0.01; *** *P* < 0.001; **** *P* < 0.0001.

## Acknowledgements

This work was supported by an ERC starting grant (716761), a Swiss National Science Foundation professorship grant (170654) to S.C.. We thank Masanori Sakaguchi for sharing the design of dual-colour miniscope, Benjamin Grewe for the calcium analysis toolset, the members of the Ciocchi laboratory for discussions and assistance with the project, Michael Känzig for genotyping the mice and Christian Dellenbach for technical assistance.

## Contributions

J.M.D., R.N. and S.C. conceived the project and designed experiments. J.M.D. performed the experiments and collected data with the help of R.N. and K.L.. J.M.D. and R.N. analysed data with help from S.C. and K.L.. S.C., R.N., J.M.D., and K.L. contributed to the writing of the manuscript. S.C. acquired funding and S.C. and R.N. supervised the project.

## Lead contact

Further information and requests may be directed to the Lead Contact, Stéphane Ciocchi.

## Materials availability

This study did not generate new unique reagents

## Data and code availability

Any original code or additional information required to reanalyse the data reported in this paper are available upon request.

## Supplementary figures

**Supplementary Figure 1.**
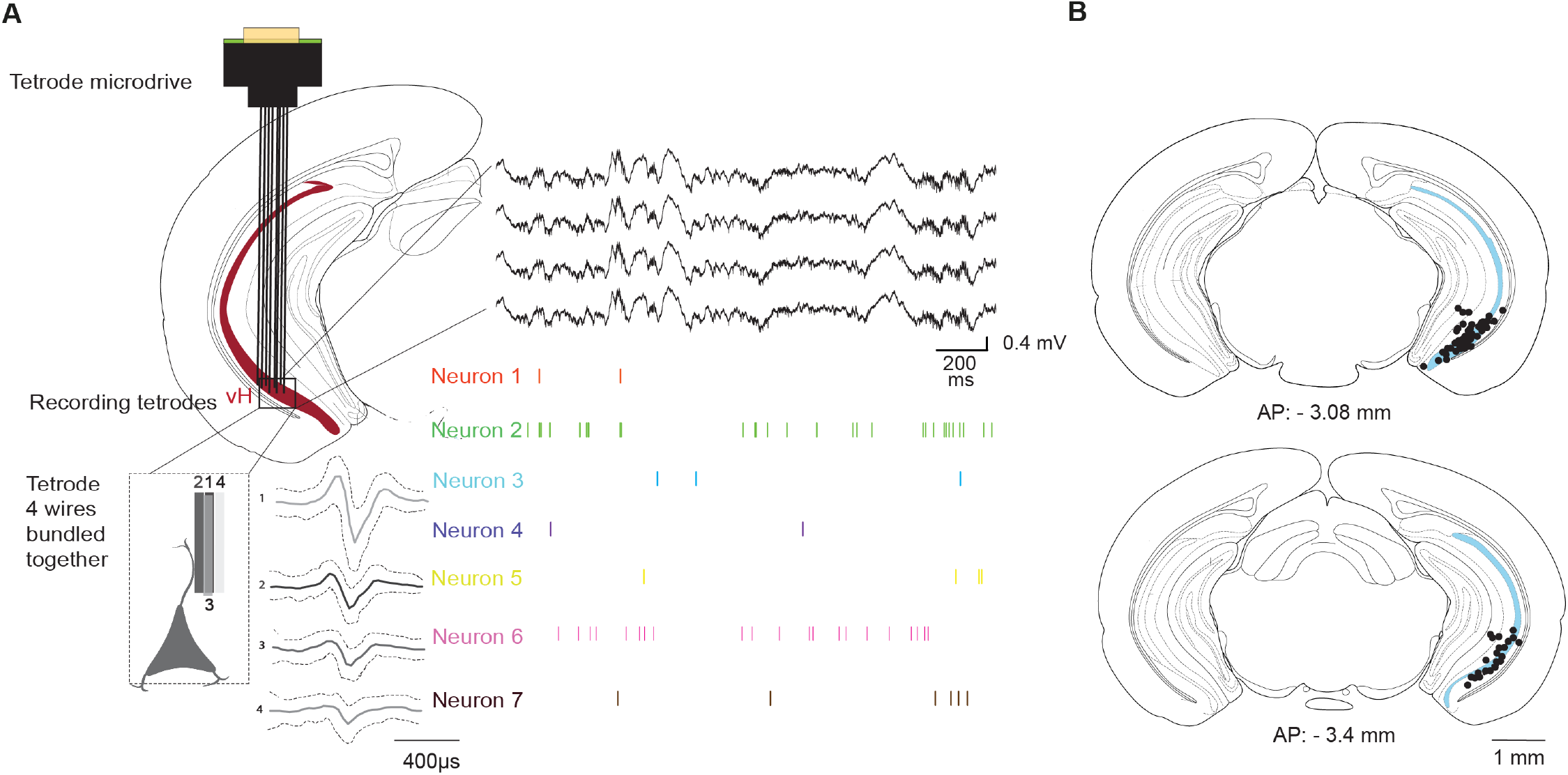
Single-unit recordings in the vH of behaving mice. **(A)** Schema of single-unit recordings using eight independently movable tetrodes in freely-behaving mice (left). Right, wide-band local field potential signal and 7 units recorded from one tetrode (4 channels). **(B)** Coronal sections of the vH showing the location of the recording sites in the vH. Each dot represents a recovered electrolytic lesion from the recorded mice (n = 23).

**Supplementary Figure 2.**
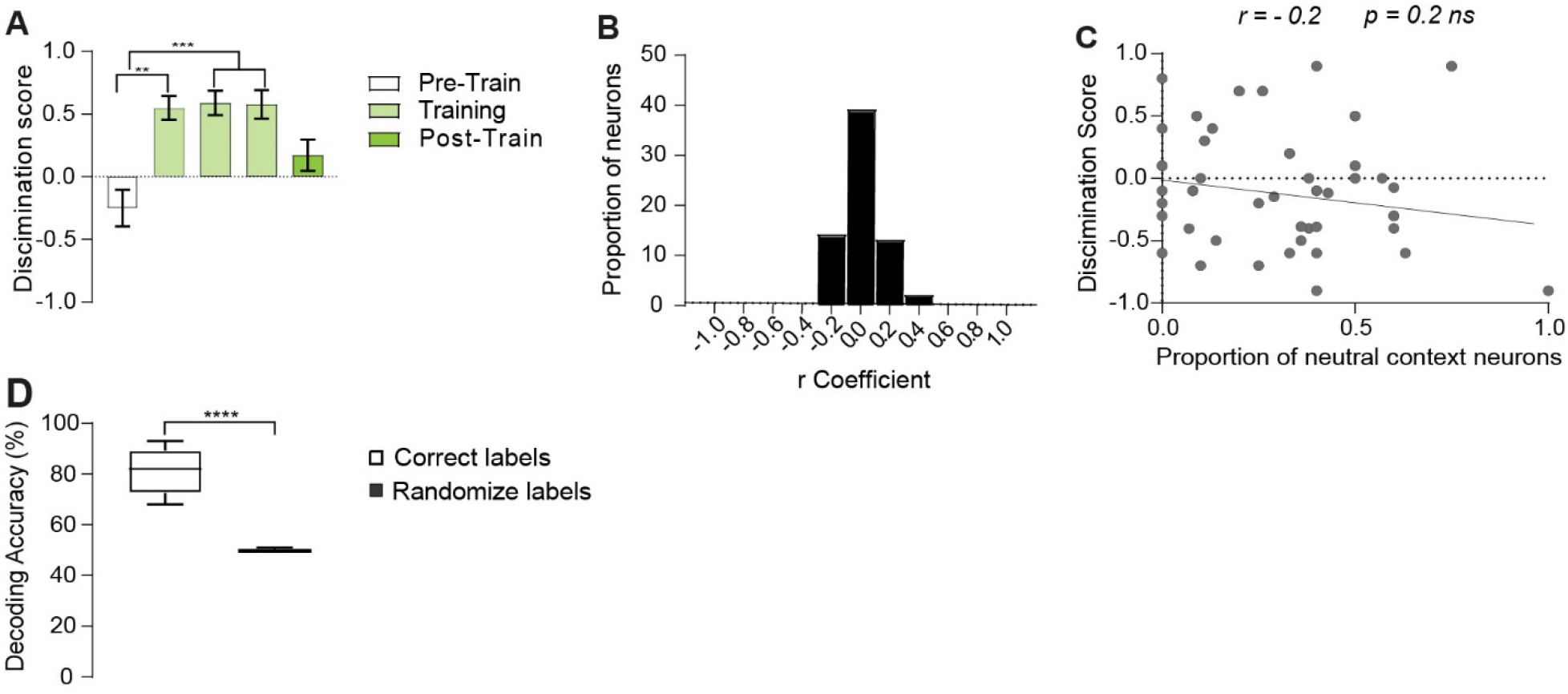
vH activity during the social reward CPP paradigm. **(A)** Discrimination score based on the difference in time spent in the social reward and neutral zones in each session of the social reward CPP. Friedman test comparison between groups, Pre-Train vs. Training 1, *P* = 0.0012, Pre-Train vs. Training 2, *P* = 0.0006, Pre-Train vs. Training 3, *P* = 0.0010, *n* = 19 mice. **(B)** Correlational analysis of firing rates vs. speed in social reward context neurons during the social reward CPP, *n* = 68 neurons, *n* = 19 mice. **(C)** Correlational analysis between behavioural social reward CPP expression during Pre-Train and Post-Train sessions in each animal and the proportion of neutral context neurons. Pearson’s r correlation, *P* = 0.1923, *n* = 23 mice. **(D)** SVM context decoding accuracy using all neurons recorded in individual mice. Unpaired t-test between groups *t*_(10)_ = 8.338, *P* < 0.0001, *n* = 94 neurons, *n* = 6 mice.

**Supplementary Figure 3.**
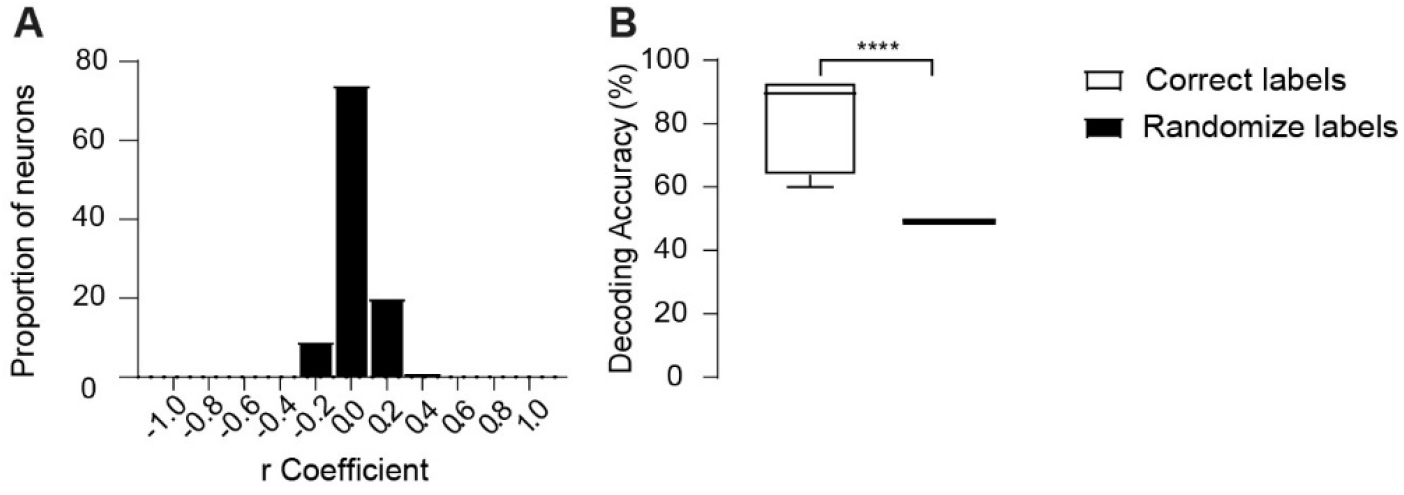
vH activity during the contextual fear conditioning paradigm. **(A)** Correlational analysis of firing rates versus speed in vH, *n* = 104 neurons, *n* = 20 mice. **(B)** SVM context decoding accuracy using all neurons recorded in individual mice. Unpaired-t test between groups *t*_(10)_ = 5.263, *P* = 0.0004, *n* = 73 neurons, *n* = 6 mice.

**Supplementary Figure 4.**
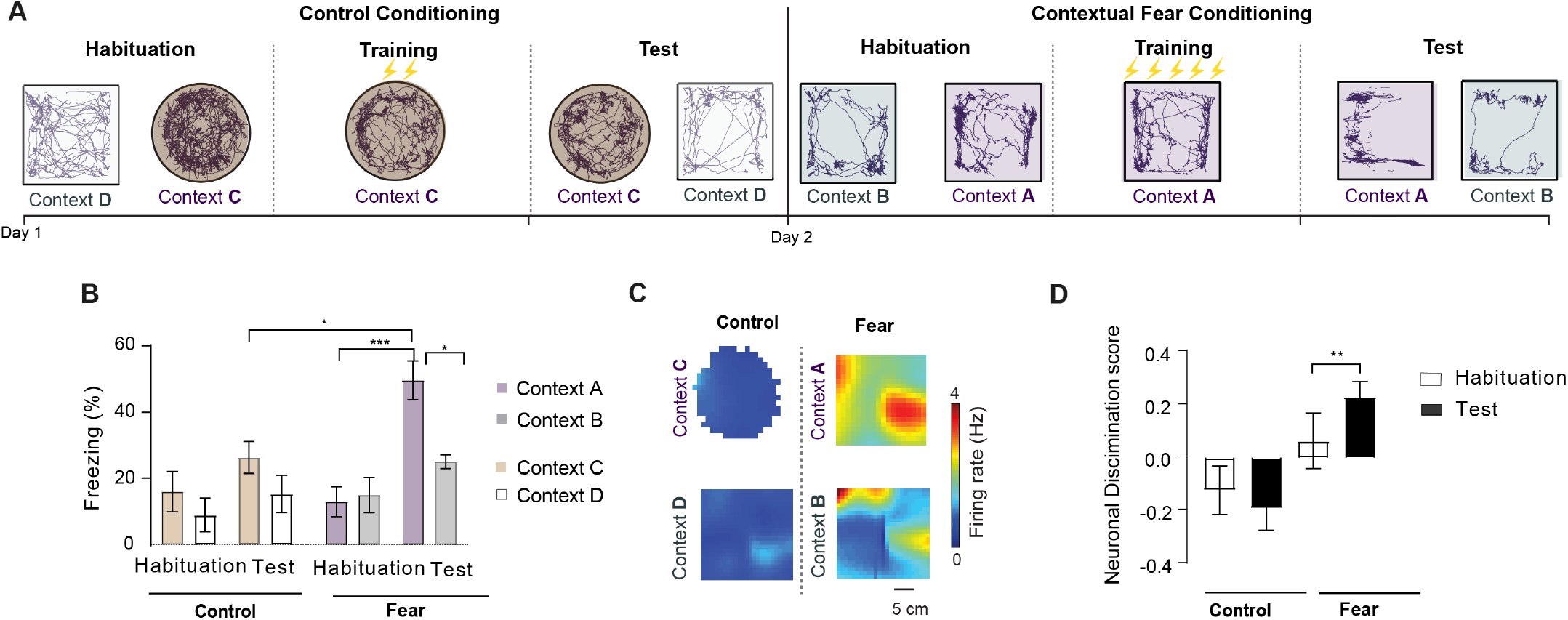
Fear context neurons are recruited upon fear learning. **(A)** Experimental design of the control ‘conditioning’ and contextual fear conditioning paradigm with representative mouse trajectories during the habituation, training, and test sessions. **(B)** Freezing levels during habituation and test sessions of the control ‘conditioning’ and contextual fear conditioning paradigm. Two-way repeated measures ANOVA between groups, Task x session interaction *F*_(3,12)_ = 2.734, *P* = 0.090. Multiple comparisons between context A vs. context C during test sessions, *P* = 0.0189, Habituation vs. Test at context A, *P* = 0.0008, Context A vs. Context B during test, *P* = 0.0172, *n* = 3 mice. **(C)** Representative examples of spatial firing rate maps of a fear context neuron during control ‘conditioning’ and contextual fear conditioning in contexts A, B, C, D. **(D)** Neuronal discrimination score activity during control ‘conditioning’ and contextual fear conditioning. Two-way repeated measures ANOVA between groups, Task x session *F*_(1,11)_ = 7.818, *P* = 0.0174. Multiple comparisons between habituation and test during fear task, *P* = 0.0089, *n* = 10 neurons out of a total of 19 recorded neurons, *n* = 3 mice.

**Supplementary Figure 5.**
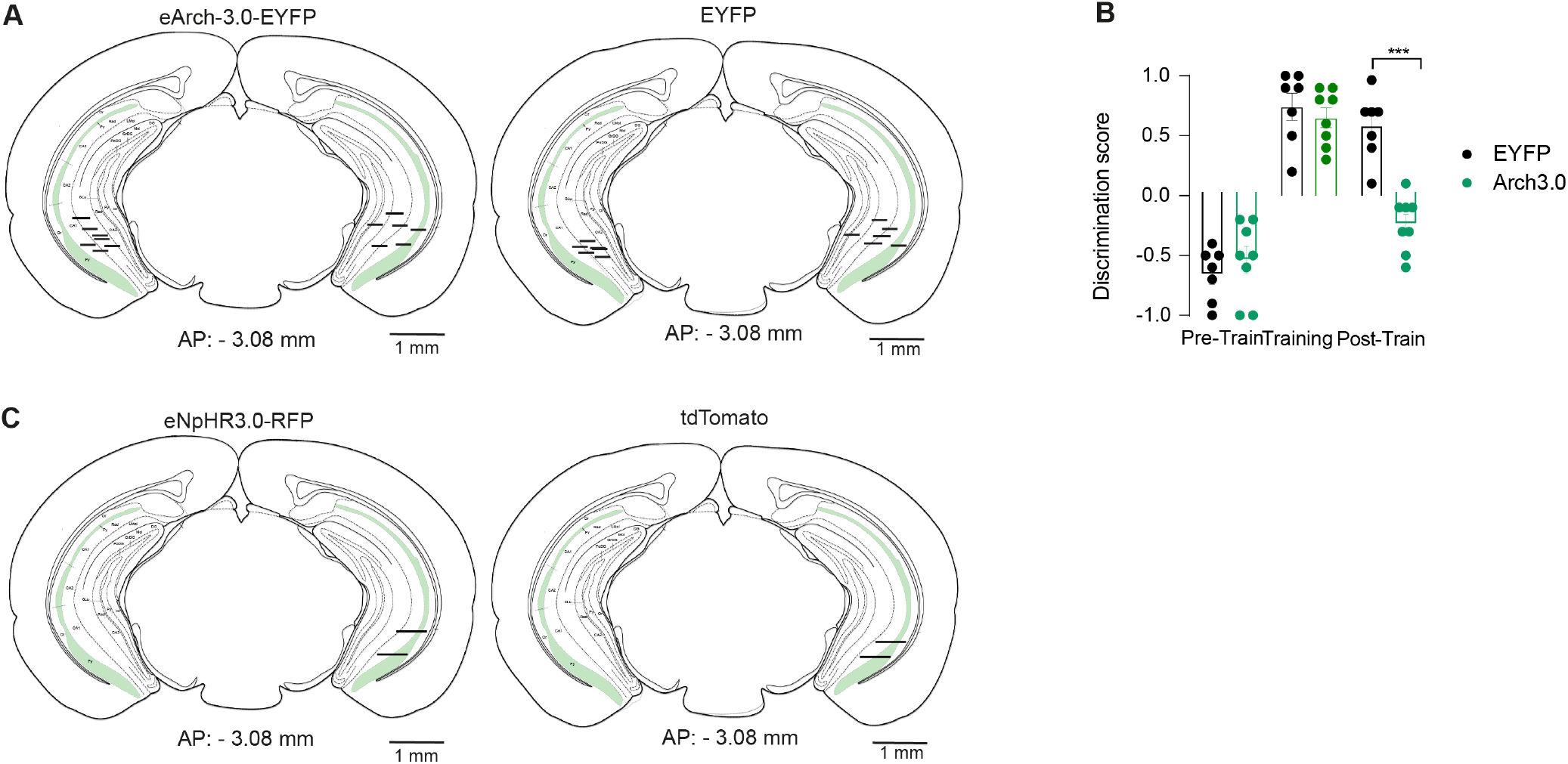
Histology of optogenetic and miniscope experiments, and behaviour-related to the social reward zone. **(A)** For each experimental group, the location of the optic fibres for each mouse is represented by a line that is mapped onto the mouse brain atlas, *n* = 7 EYFP mice, *n* = 8 Arch mice. The histology of one Arch and one EYFP mouse (bilateral vH) and another Arch mouse (unilateral vH) could not be properly recovered due to slicing issues and are thus not shown. **(B)** Discrimination score based on the time spent in the social reward zone. Two-way repeated measures ANOVA between groups. Condition vs. session interaction *F*_(2,26)_ = 3.421, *P* = 0.0480. Multiple comparisons between EYFP vs. eArch3.0 during Post-Train session, *P* = 0.0002, *n* = 7 EYFP, *n* = 8 eArch3.0 mice. **(C)** The GRIN lens location for each mouse of each experimental group is represented by a line and that is mapped onto a mouse brain atlas (*n* = 2 tdTomato mice, *n* = 2 eNpHR3.0 mice).

**Supplementary Figure 6.**
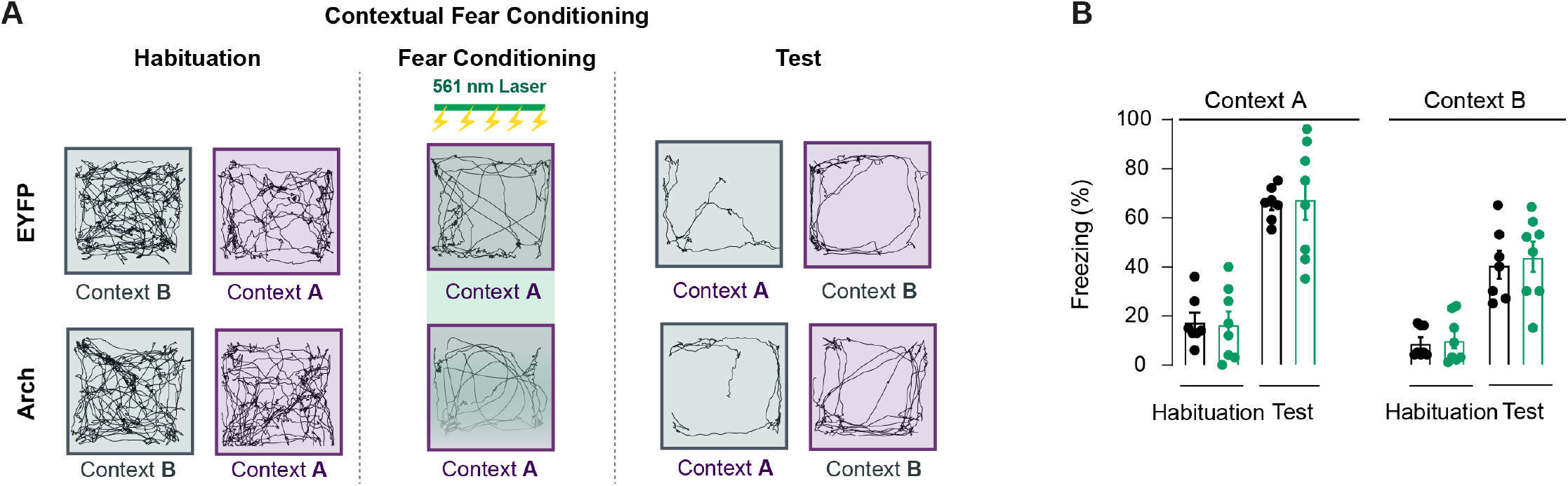
Contextual fear memory does not require locus coeruleus projections to the vH. **(A)** Schema of the optogenetic inhibition protocol of LC projections to the vH during contextual fear conditioning with representative mouse trajectories. **(B)** Corresponding freezing levels during contextual fear conditioning. Condition vs. session interaction *F*_(3,36)_ = 0.07331, *P* = 0.09739, *n* = 7 EYFP, *n* = 8 eArch3.0 mice.

